# A circRNA-based uricase replacement therapy for sustained treatment of hyperuricemia

**DOI:** 10.64898/2026.03.19.712815

**Authors:** Zheyu Zhang, Jiewen Zhong, Kai Zhang, Jiang Hu, Yun Yang, Zefeng Wang

## Abstract

Hyperuricemia, a major risk factor for gout and kidney disease, arises from the evolutionary loss of human uricase and remains a significant medical challenge due to its high prevalence. However, limited therapeutic options are available for refractory hyperuricemia that typically require long-term treatment. Here we developed a circRNA-based uricase replacement strategy and evaluated its efficacy in uricase-knockout mice as a model for severe hyperuricemia. Lipid nanoparticle-mediated delivery of circRNA enabled efficient in vivo expression of an engineered human-like uricase, which rapidly reduced serum urate levels after a single injection and maintained the urate-lowering effect for up to 10 days. Repeated administration led to sustained urate reduction for 10 weeks, mitigated renal injury, and exhibited favorable biosafety. These findings highlight the therapeutic potential of circRNA-based uricase replacement for the long-term treatment of hyperuricemia and its associated complications.

## Introduction

Hyperuricemia, primarily resulted from impaired urate excretion, is clinically defined as a serum urate concentration above certain threshold (typically >420 μmol/L in men and >360 μmol/L in women) (*1, 2*). As a poorly soluble end-product of purine catabolism (*3*), uric acid is efficiently degraded in most mammals by uricase (UOX, urate oxidase), which catalyzes the oxidation of urate to 5-hydroxyisourate (*4*), a metabolite further processed into highly soluble allantoin, urea, or ammonia for excretion (*5*). However, the uricase activity gradually declined during human evolution and was completely lost approximately 15-20 million years ago, leaving only a pseudogene in the genome (*6, 7*). Although high urate levels may have conferred selective advantages to early hominoids against food shortage (*7, 8*), hyperuricemia has now emerged as a significant risk factor for a range of pathological conditions in the current human population that has an improved living standards and longer life expectancy (*1, 9, 10*). Notably, hyperuricemia is the most important cause of gout (*9, 11*), in which monosodium urate (MSU) crystals deposit in peripheral joints, resulting in the activation of the NLRP3 inflammasome that triggers gout flares with the release of interleukin-1β (*12, 13*). Similar MSU deposition within renal tissues also contributes to chronic kidney disease (CKD) (*14, 15*).

Beyond its direct role in gout and CKD, emerging evidence has also linked hyperuricemia to cardiovascular diseases, metabolic syndrome, hypertension, and diabetes (*1, 16–18*). Importantly, hyperuricemia is increasingly recognized as a disease-modifying factor rather than a purely symptomatic condition. Recent studies have demonstrated that sustained urate reduction significantly delays CKD progression and reduces both all-cause and cardiovascular mortality (*19*). Therefore, effective urate-lowering therapy is clinically meaningful even in individuals with asymptomatic hyperuricemia. Additionally, during the treatment of cancers, massive tumor cell lysis following chemotherapy or radiotherapy can result in tumor lysis syndrome (TLS), leading to a sudden and excessive accumulation of urate that exceeds homeostatic control to cause acute systemic toxicity and kidney injury, necessitating rapid urate elimination rather than only suppression of urate production (*20*). Globally, hyperuricemia affects 2.6-36% of individuals across different populations and regions, with prevalence continuing to rise (*1, 21*). Because hyperuricemia can remain asymptomatic for long periods in certain populations (*22*), the true prevalence is likely underestimated.

Despite the need for long-term management of hyperuricemia and its large patient population, current therapeutic options remain limited. Allopurinol, a first-line xanthine oxidase inhibitor, effectively reduces *in vivo* urate synthesis but carries the risk of severe cutaneous adverse reactions, particularly in patients with the HLA-B*58:01 allele that is common in Asian populations (*23–26*). Meanwhile, it cannot rapidly catabolize pre-existing urate, limiting its application in TLS. Another xanthine oxidase inhibitor, febuxostat, has been associated with increased cardiovascular risk and mortality (*27, 28*). In addition, inhibition of xanthine oxidase can elevate xanthine and hypoxanthine levels, potentially causing kidney xanthine stones that necessitate careful dose adjustment and maintenance of adequate urine flow (*29*). Beyond xanthine oxidase inhibitors, benzbromarone and probenecid function as uricosuric agents to promote urate excretion; however, they are associated with rare but notable hepatotoxicity and nephrotoxicity, respectively (*9*). Rasburicase and pegloticase are recombinant uricases that provide an alternative strategy by directly degrading urate. Yet, the use of rasburicase in TLS is restricted by considerable adverse effects (*30*) and its short half-life (*31–33*). Although the pegloticase extends the half-life *via* PEG modification, this drug still suffers from high immunogenicity with approximately 26% of patients in phase III trials experiencing infusion-related reactions primarily due to anti-PEG antibodies and loss of urate-lowering efficacy (*34, 35*).

Given these limitations, novel therapeutic strategies for durable and safe urate control are urgently needed. Following the success of two mRNA vaccines against COVID-19 (*36, 37*), RNA-based therapeutics have attracted increasing attention (*38–40*). Unlike conventional gene-editing approaches, RNA-based therapies do not result in permanent genomic integration, thereby offering an improved safety profile (*38, 40*). Moreover, compared with traditional protein replacement strategies, it enables direct intracellular expression of the target protein to exert its natural function, circumventing the requirement for the complex and expensive *in vitro* protein production and purification (*41, 42*). Circular RNA (circRNA) is a single-stranded RNA with covalently closed structure (i.e., lacking free 5’ and 3’ ends). Recent studies have shown that circRNAs containing internal ribosome entry sites (IRES) can drive efficient cap-independent translation *in vivo* (*43–45*), while exhibiting superior stability due to their resistance to RNase R–mediated degradation (*46*). Moreover, *in vitro* synthesized circRNA of sufficient purity exhibits low innate immunogenicity without nucleotide modifications, which are typically required for linear mRNA drugs (*47–49*). These properties highlight the potential of circRNA as a promising therapeutic modality for long-term protein replacement that require sustained protein expression, repeated dosing, improved biosafety and reduced manufacturing complexity.

In this study, we demonstrate the long-term therapeutic potential of repeated circRNA-based therapy in a uricase knockout mouse model of severe hyperuricemia. We delivered a circRNA encoding a humanized uricase *via* lipid nanoparticles (LNPs), aiming to restore hepatic uricase expression and reduce serum urate levels. We systematically assessed therapeutic efficacy across a range of doses and dosing intervals, and conducted preliminary evaluations for the biosafety profile. The results suggest a long-term efficacy of circRNA therapy in reducing serum urate and kidney damages, demonstrating its potential as a novel treatment for refractory hyperuricemia.

## Results

### Efficient expression of engineered UOX circRNAs in human cells

To mimic the clinical application of exogenous uricase in human hyperuricemia, three artificial engineered UOX variants were selected for initial testing: rHU (*50*), reconstructed from the human UOX pseudogene; PBC (*34*), the pig–baboon chimeric UOX sequence of the clinical drug pegloticase; and An96 (*8*), identified through phylogenetic analysis of uricase. All three UOX variants have been reported to exhibit strong uricase activity, high stability, and promising biosafety (*8, 34, 50*).

The optimized coding sequences of three UOX variants were inserted into circRNAs containing the Coxsackievirus B3 (CVB3) IRES that can drive efficient translation (*51*). An N-terminal FLAG tag sequence was incorporated for detection, while the native C-terminal peroxisomal targeting signal sequence was retained (Fig. 1A). The DNA template is transcribed by T7 RNA polymerase, and the linear RNA precursor is circularized during *in vitro* transcription by separated self-splicing introns at both ends of the DNA template (Fig.1A and Fig. 1B). Three UOX circRNAs were synthesized and purified according to previously established methods (*49*).

**Figure 1.**
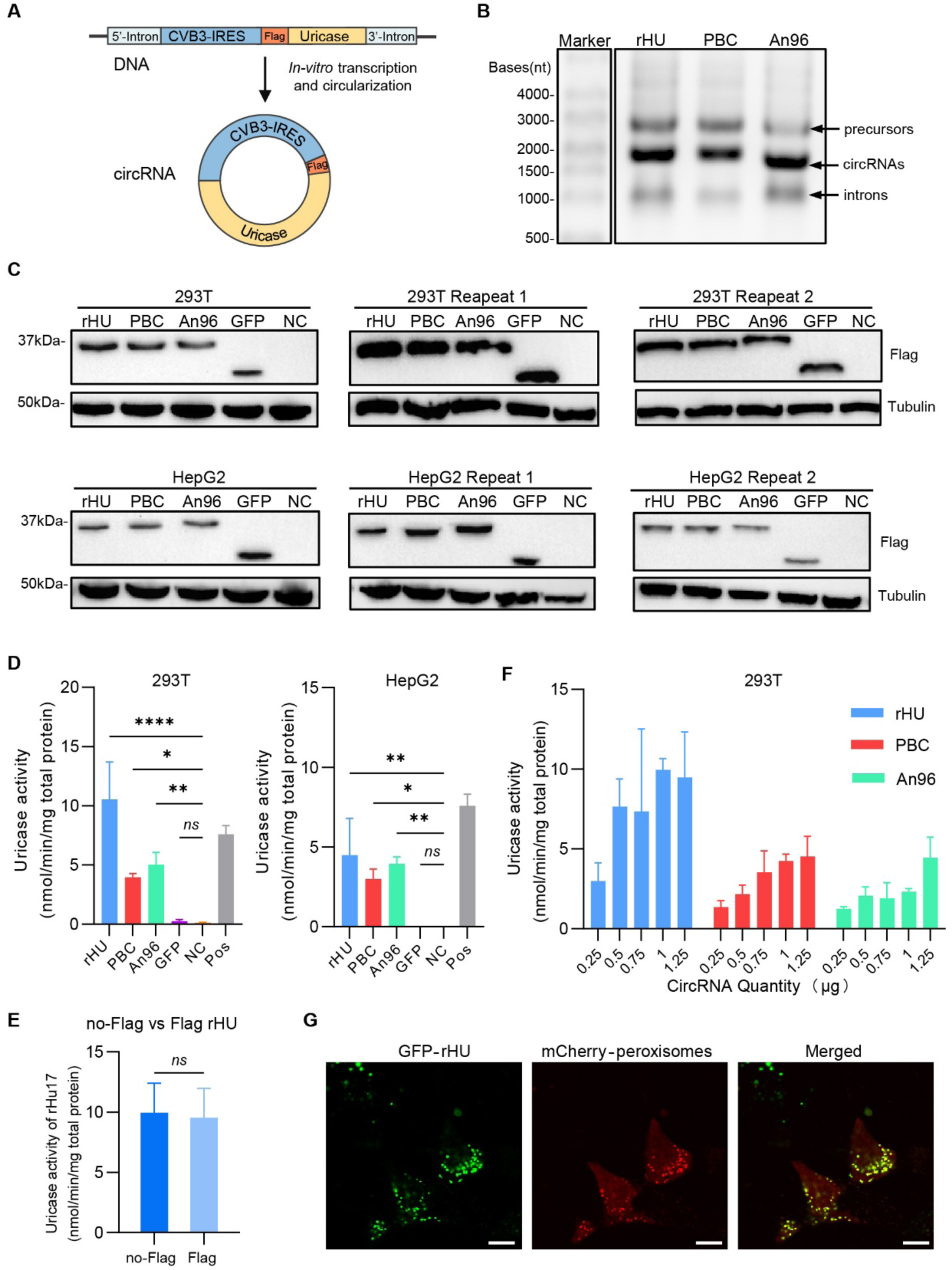
Characterization of UOX circRNA synthesis and expression. (A) Design of the UOX circRNAs. Each circRNA comprises two separated parts of group II self-splicing introns, a CVB3 IRES, a FLAG tag and the uricase coding sequence. CircRNAs were synthesized from DNA templates by *in vitro* transcription (IVT), during which the group II intron parts flank the linear RNA precursors to enable autocatalytic circularization. The CVB3 IRES drives cap-independent translation, and the FLAG tag was used for detection of uricase. (B) Gel electrophoresis of IVT products from three UOX variants (rHU, PBC, An96). Each IVT reaction produced linear RNA precursors, circRNAs, and excised introns. (C) Western blot analysis of 293T and HepG2 cell lysates transfected with rHU, PBC, An96 or GFP circRNAs. Lysates from untransfected cells served as the negative control (NC). Three independent transfections were performed on separate days, and all replicates are shown. Proteins translated from circRNAs were detected using an anti-FLAG antibody and normalized to α-tubulin. (D) Uricase activity assays of 293T and HepG2 cell lysates transfected with rHU, PBC, An96 or GFP circRNAs (8µg total protein per sample). Cell lysates from untransfected cells served as the negative control (NC). The uricase (0.02µg, corresponding to 0.25% of total lysate proteins, commercially purchased) was used as the positive control (Pos). Lysates or the uricase were incubated with 0.125 mM uric acid in 0.1 M sodium borate buffer (pH 8.5) at 37°C for 120 min, and the uricase activity was quantified by the decrease in absorbance at 293 nm resulting from uric acid oxidation. Data were presented as mean ± SD. Statistical significance was determined by one-way ANOVA with Dunnett’s multiple comparison test. P < 0.05 was considered significant (*ns* > 0.05, \**p* < 0.05, \*\**p* < 0.01, \*\*\**p* < 0.001, \*\*\*\**p* < 0.0001). (E) Comparison of enzymatic activity of 293T cell lysates transfected with rHU circRNA with or without a FLAG tag. Data were presented as mean ± SD. Statistical significance was assessed by unpaired t-test; *P* > 0.05 was considered not significant (*ns*). (F) Dose-dependent uricase activity of 293T cell lysates transfected with increasing amounts of UOX circRNAs (0.25-1.25 µg) encoding rHU, PBC and An96. (G) Intracellular fluorescence co-localization of GFP-fused rHU (green) and mCherry fused with the peroxisomal targeting sequence (C-terminal Ser-Arg-Leu, red).

Upon transfection to 293T and HepG2 cells, we found that all three UOX circRNAs directed translation of uricase proteins with comparable efficiency (Fig. 1C and Fig. S1). Among them, the rHU displayed the highest enzymatic activity in cell lysates (Fig. 1D), while the control group exhibited no uricase activity, consistent with their human origin and lack of the endogenous uricase (Fig. 1D). Notably, the FLAG tag did not affect rHU activity since no significant difference was observed between the lysates of the cells transfected with rHU circRNA with or without the tag (Fig. 1E). In addition, the uricase activities were positively correlated with the dose of transfected circRNAs (Fig. 1F). Fluorescence imaging further revealed that GFP-fused rHU strongly overlapped with mCherry-labeled peroxisomes (Fig. 1G), suggesting that exogenously expressed uricase in human cells was targeted to peroxisomes *via* its native targeting signal, consistent with the natural localization of uricase in other species (*52, 53*).

### Expression kinetics and biosafety of circUOX-LNP in wild type mice

To examine the expression of uricase from circRNAs in animal models, we employed the circRNA encoding UOX-rHU (circUOX) that demonstrated relatively higher efficacy in cell assays. The circUOX-LNP was formulated using the ALC-0315 formulation which is clinically approved for mRNA vaccines and showed favorable liver-targeted delivery efficiency and good biosafety performance via intravenously administration in recent research (*54*). We purified circRNA to high purity (Fig. S2A) for subsequent microfluidic encapsulation and purification, the resulting circUOX-LNPs exhibited a mean particle size of 74.3 ± 1.6 nm and an encapsulation efficiency of 89% (Fig. S2B).

The circUOX-LNPs were administered *via* tail vein injection in wild-type mice using three different doses (0.3 to 1.2mg/kg, Fig. 2A). We found that the hepatic circUOX increased in a dose-dependent manner, with markedly enhanced signals observed in the high-dose group (Fig. 2B). The circUOX was readily detectable by RT-qPCR in livers at day 1 post-injection and remained at a lower level through days 3 and 5. Using western blot analysis, we observed a clear dose-dependent increase in hepatic uricase expression, with the 0.6 mg/kg and 1.2 mg/kg groups maintaining appreciable protein levels even at day 5, indicative of a prolonged therapeutic window (Fig. 2C).

**Figure 2.**
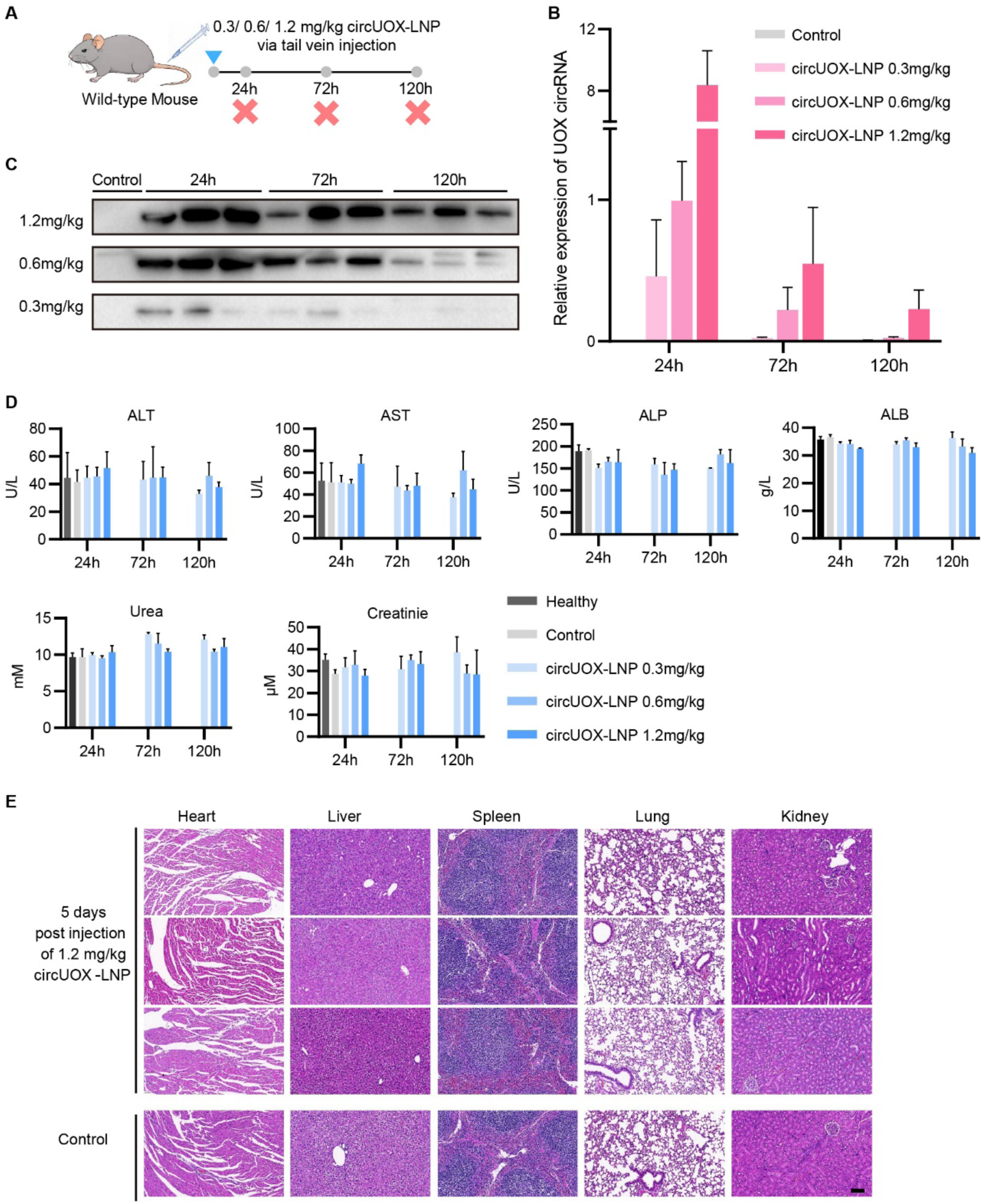
Expression of UOX (rHU) circRNA in mouse livers and evaluation of its biosafety. (A) Schematic of the experimental design for evaluating expression kinetics and biosafety in wildtype C57 mice. CircUOX-LNPs (UOX-rHU) were administered intravenously at doses of 1.2, 0.6, or 0.3 mg/kg. Mice (*n* = 3 per group) were sacrificed on days 1, 3, and 5 post-injection for serum and tissue collection. Buffer-injected mice (*n* =3) were sacrificed on day 1 and served as the control group. Uninjected mice (*n*=3) were employed as the healthy group. (B) Hepatic UOX circRNA levels on days 1, 3, and 5 following intravenous injections at doses of 1.2, 0.6, or 0.3 mg/kg. Buffer-treated mice sacrificed on day 1 served as controls. RT-qPCR was performed using primers flanking the back-splicing junction, and circRNA levels were normalized to GAPDH mRNA levels. Data were presented as mean ± SD. (C) Western blot analysis of exogenous uricase rHU in liver lysates on days 1, 3, and 5 following intravenous injections at doses of 1.2, 0.6, or 0.3 mg/kg. Three buffer-treated mice sacrificed on day 1 served as controls. Proteins immunoprecipitated from 2 mg of total lysate protein using anti-FLAG magnetic beads were used for the Western blot. (D) Serum biochemical indicators for each group. ALT, AST, ALP and ALB were measured to assess hepatic function, while urea and creatinine were measured to assess renal function. Data were presented as mean ± SD. No statistical significance was determined by one-way ANOVA (*p* > 0.05). (E) Hematoxylin and eosin (H&E) staining of hearts, livers, spleens, lungs, and kidneys from control mice and mice 5 days after a single 1.2 mg/kg dose of circUOX-LNPs. Scale bar, 100 µm.

To evaluate hepatic and renal toxicity, biochemical indicators of liver and kidney function were measured, including alanine aminotransferase (ALT), aspartate aminotransferase (AST), alkaline phosphatase (ALP), albumin (ALB), urea and creatinine (Crea). The results revealed no significant abnormalities across all groups (Fig. 2D), suggesting that even the high-dose circUOX-LNP was well tolerated without causing significant adverse effects on liver or kidney function. Consistently, histological examination of major organs including hearts, livers, spleens, lungs, and kidneys showed no obvious pathological changes (Fig. 2E), further supporting the biosafety of the circUOX-LNP.

### Urate reduction and renal protection by circUOX-LNP treatment

To evaluate the therapeutic efficacy of circUOX-LNP, we employed the uricase-knockout (Uox⁻/⁻) mice as a model of severe hyperuricemia (*55*) and treated the mice with circUOX-LNP at various dosing schedules. A single injection rapidly reduced the serum urate levels below saturation threshold (∼400 µM) within 24 h in a dose-dependent manner, with the urate-lowering effect lasting up to 10 days (Fig. 3A). In contrast, the control mice injected with buffers maintained persistently high urate levels above the saturation threshold. The repeated dosing further demonstrated a prolonged efficacy: each circRNA-LNP injection triggered a rapid and robust reduction in serum urate to near-normal levels (Fig. 3B). Following the third dose on day 8, urate levels were significantly suppressed for another 7 days and stayed below baseline through day 20 (Fig. 3B).

**Figure 3.**
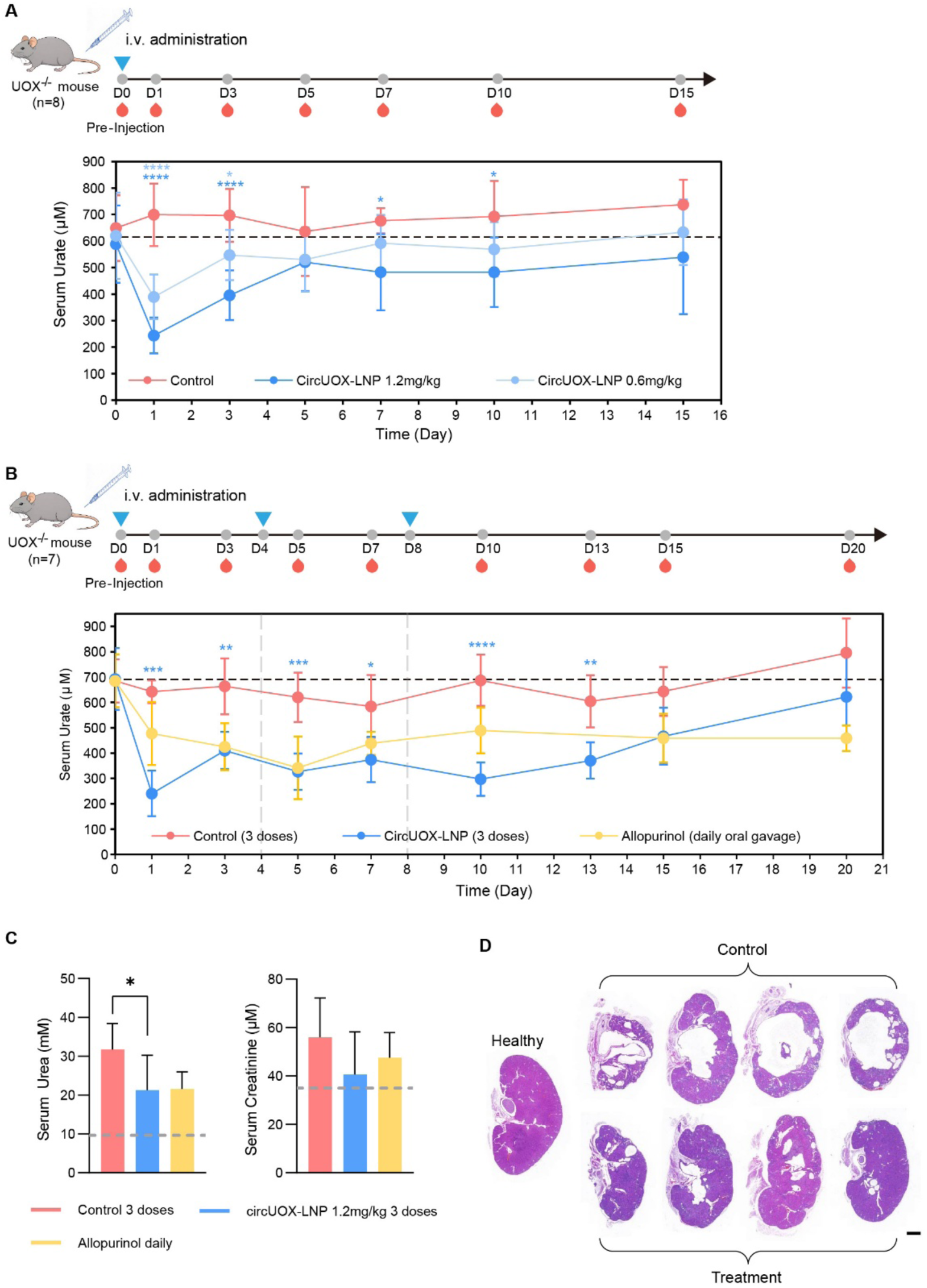
Uric acid-lowering effects of circUOX-LNP. (A) Urate-lowering effects in 6-week-old UOX^⁻/⁻^ mice (*n* = 8) following a single injection of circUOX-LNP at two doses (0.6mg/kg and 1.2mg/kg) or an equal volume of buffer solution as a control. Serums were collected from eye sockets for urate assays at each time point (days 1,3,5,7,10,15 post-injection). (B) Urate-lowering effects in 8-week-old UOX^⁻/⁻^ mice (*n* = 7) following three injections of 1.2mg/kg circUOX-LNP or an equal volume of buffer as a control. Injections were administered every 4 days. Allopurinol-treated UOX^⁻/⁻^ mice (*n* = 3) received daily oral gavage at 15 mg/kg, corresponding to the clinical starting dose in humans, and served as a positive control. Serums were collected for urate assays at days 1,3,5,7,10,13,15 and 20 after the first injection. Data in (A-B) were presented as mean ± SD. P value was calculated by two-way ANOVA with Bonferroni’s multiple comparisons test. *P* < 0.05 were considered statistically significant (\**p* < 0.05, \*\**p* < 0.01, \*\*\**p* < 0.001, \*\*\*\**p* < 0.0001). (C) Serum renal function indicators (urea and creatinine) in UOX^⁻/⁻^ mice at the end of the experiments (day 20). Data for the healthy group were obtained from wild-type C57 mice with no injection (Fig. 2D). Gray dashed lines indicate the urea and creatinine levels of healthy wild-type C57 mice (data from Fig. 2D). Data were presented as mean ± SD. Statistical significance was determined by one-way ANOVA with Dunnett’s multiple comparison test. P < 0.05 was considered significant (*ns* > 0.05, \**p* < 0.05, \*\**p* < 0.01, \*\*\**p* < 0.001, \*\*\*\**p* < 0.0001). (D) H&E staining of UOX^⁻/⁻^ mice kidneys from the 3-dose circUOX-LNP-treated group and the 3-dose buffer-treated control group. The kidney from an un-injected wild type mouse were used as a healthy control. Scale bar, 1mm.

Compared with the control group that received daily oral gavage of allopurinol (15 mg/kg, a dose corresponding to the clinical starting regime), circUOX-LNP treatment induced a more rapid urate reduction starting from day 1 (Fig. 3B). Conversely, allopurinol required approximately 3-5 days to reach urate levels comparable to those in the circUOX-LNP group, likely due to slower excretion of pre-existing urate compared with the direct urate degradation.

The untreated mice exhibited an elevated level of serum urea and creatinine, reflecting renal damage. We found that the mice with 3-dose circUOX-LNP treatment significantly reduced the serum urea level, indicative of partial relief of renal injury (Fig. 3C). The creatinine levels were also reduced by ∼30%, although such change is not statistically significant due to the small sample size. Consistently, the untreated mice showed pronounced hydronephrotic dilatation accompanied by a marked loss of renal parenchyma, suggesting extensive renal damage (Fig. 3D). However, the mice treated with circUOX-LNP showed significant reduction of hydronephrosis, suggesting a protective effect against hyperuricemia-induced renal injury (Fig. 3D).

To evaluate the biosafety of circUOX-LNP in UOX^-/-^ mice, we collected the serum and major tissues at the end of the experiment (day 20) for measurements. Several biochemical indicators of liver function still remained in normal ranges after the 3-dose treatment (Fig. S3A). In addition, the histology examination of major organs of 3-dose treatment group including hearts, lungs, spleens and livers showed normal morphology similar to the untreated group (Fig. S3B), suggesting that the repeated circRNA-LNP administration did not produce obvious drug-related adverse effects.

### Reversal of transcriptome alteration caused by hyperuricemia

To further assess whether circUOX-LNP treatment could alleviate the transcriptomic changes linked to renal injury, we next performed RNA-seq analysis of mouse kidneys. The kidneys of UOX^-/-^ mice (i.e., diseased) showed remarkable transcriptomic alterations relative to wild-type healthy controls (|fold change| > 1.5, adjusted *p* < 0.05), whereas circUOX-LNP treatment partially restored normal expression in the 388 upregulated and 350 downregulated genes (Fig. 4A). This shift toward normal expression patterns is consistent with the alleviation of renal injury. Functional enrichment analyses using Metascape also indicated that the restored upregulated genes were enriched in inflammatory response pathways including NF-kappa B, IL-17 and TNF signaling pathway, consistent with reduced Inflammatory response and renal injury (Fig. 4B). Meanwhile, the restored downregulated genes were enriched in amino acid metabolism, small-molecule transport, and circadian rhythm, suggesting a potential role in the recovery of normal renal functions (Fig. 4B).

**Figure 4.**
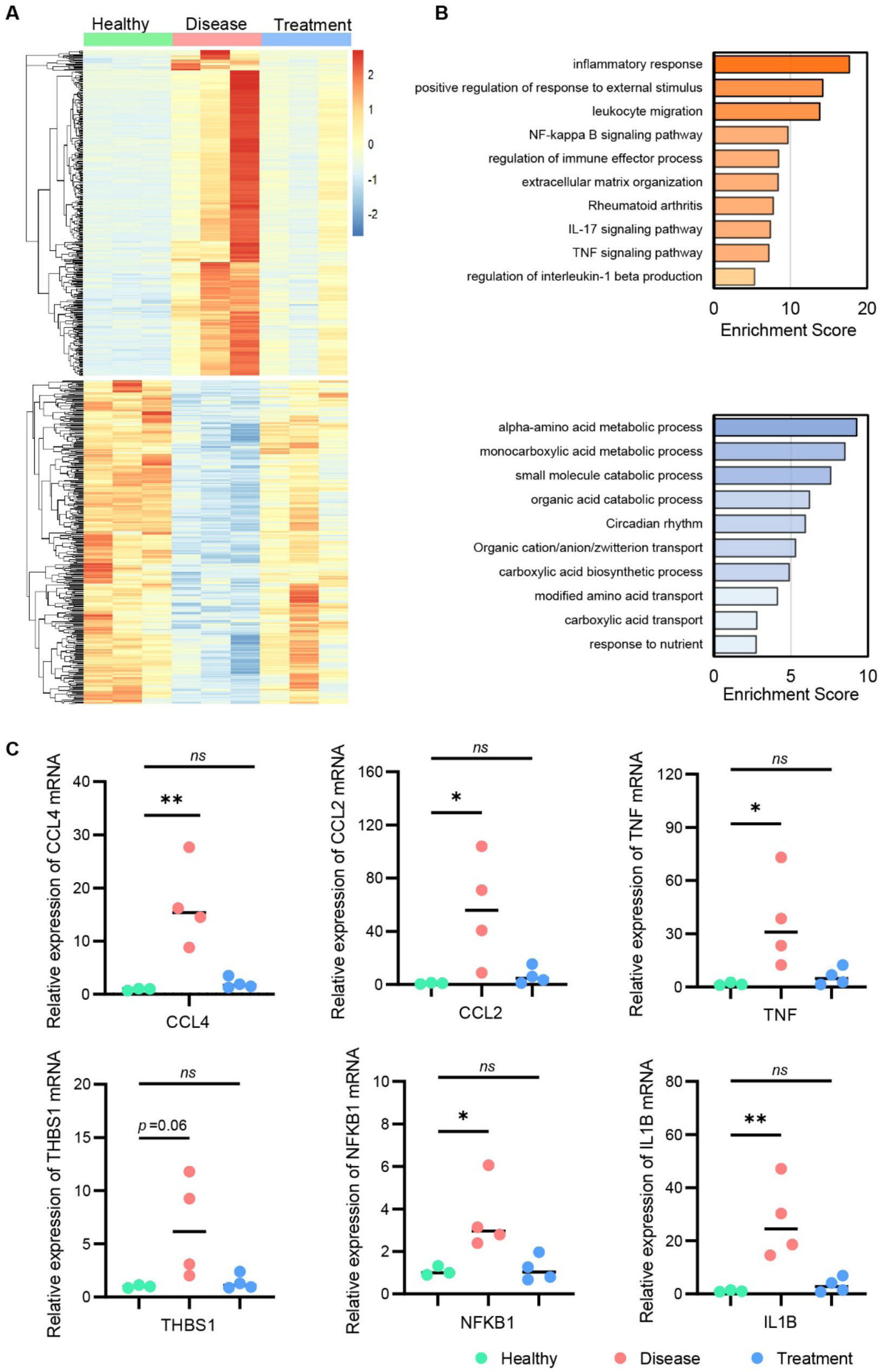
Renal protective effect of circUOX-LNP. (A) Heatmap illustrating the reversal of transcriptomic alternations in diseased kidneys after treatment. Wild type C57 mice served as the healthy group, and the hyperuricemia UOX^-/-^ mice treated with three injections of 1.2mg/kg circUOX-LNP or buffer served as treatment and control group. Data were presented as mean FPKM values, z-score normalized (*n* = 3 per group). (B) Functional enrichment analysis of restored genes after treatment, performed using Metascape. Enriched terms for restored upregulated genes are shown in red, and those for downregulated genes are shown in blue. (C) Relative expression levels of restored genes in RNA-seq of mouse kidneys from each group (*n* = 3 in the healthy group, *n* = 4 in the treatment and control groups). RT-qPCR was performed for CCL4, CCL2, TNF, THBS1, NFKB1 and IL1B, normalized to GAPDH mRNA levels. The lines in charts represent median values. P value was calculated by one-way ANOVA with Dunnett’s multiple comparison test. *P* < 0.05 were considered statistically significant (*ns* > 0.05, \**p* < 0.05, \*\**p* < 0.01).

To validate the sequencing results, RT-qPCR was performed for CCL4, CCL2, TNF, THBS1, NFKB1, and IL1B (Fig. 4C). Among these genes, TNF and IL1B encode pro-inflammatory cytokines that activate inflammation-associated pathways, including the NF-κB signaling pathway (*56*). CCL2 and CCL4 act as inflammatory chemokines, and notably, CCL2 has been proved implicated in renal injury (*57*). In addition, THBS1 overexpression is associated with renal inflammation and fibrosis (*58, 59*). All these 6 upregulated genes in the diseased kidneys exhibited a significant or marginally significant decrease following circRNA-LNP treatment, confirming a transcriptomic attenuation of renal injury in treated mice (Fig. 4C).

### CircUOX-LNP as a long-term urate-lowering therapy in mouse model

To simulate the long-term therapy in human hyperuricemia, we evaluated the urate-lowering efficacy of circUOX-LNP administered once every eight days. In previous studies, the UOX^⁻/⁻^ mice tend to develop severe kidney injury during prolonged hyperuricemia (Fig. 3D). Therefore, younger mice (4 weeks old) were used for the long-term treatment experiment, allowing urate lowering before extensive renal damage occurred.

Serum urate monitoring demonstrated that circUOX-LNP produced a sustained and potent urate-lowering effect throughout the treatment period (Fig. 5A), whereas untreated controls showed progressive increase in serum urate levels. In addition, each administration reduced urate levels for approximately one week without attenuation (Fig. 5A), confirming the efficacy of repetitive dosing.

**Figure 5.**
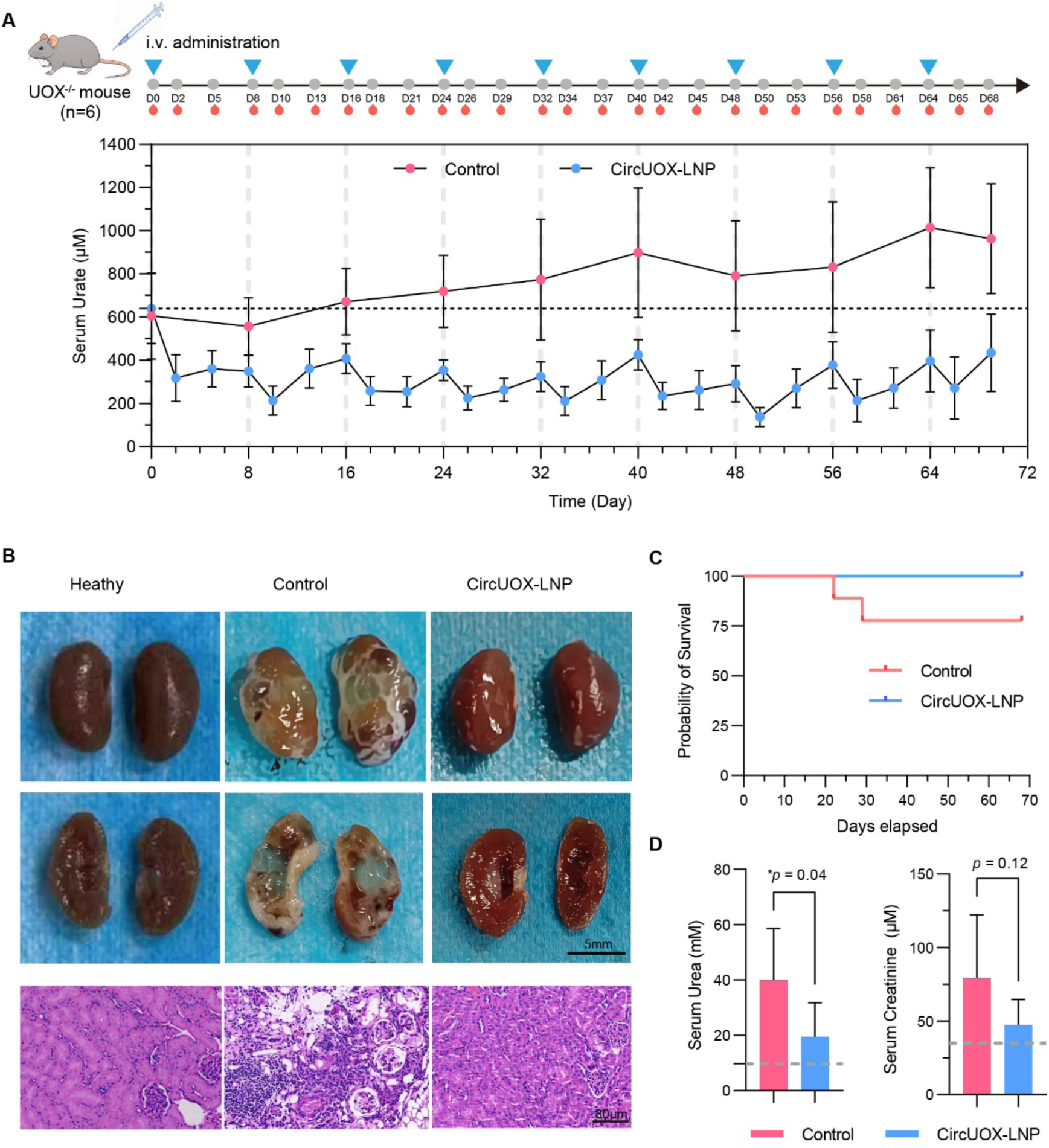
Long-term efficacy of circUOX-LNP in hyperuricemia mouse. (A) Long-term urate-lowering effects in 4-week-old UOX^⁻/⁻^ mice (*n* = 6) following eight injections of 1.2mg/kg circUOX-LNP or an equal volume of control buffer. Injections were administered once every eight days. Serums were collected via retro-orbital blood collection on days 2, 5, 8 after each administration. (B) Photographs and H&E strained kidney sections from healthy mice, control mice and treatment mice. Wild type mouse served as the healthy group, whereas UOX^⁻/⁻^ mice receiving 8 injections of circUOX-LNP or buffer served as the treatment and control groups. Scale bars: 5 mm (photos) and 40 µm (H&E sections). (C) Survival curve of mice in the treatment group and control group (*n* = 6 in treatment group, *n* = 8 in control group and 2 UOX^-/-^ mice of control group died during the experiment. (D) Serum renal function indicators (urea and creatinine) at the end of the long-term experiment (Day 69). Gray dashed lines indicate the urea and creatinine levels of healthy wild-type C57 mice (data from Fig. 2D). Data were presented as mean ± SD. P value was calculated by unpaired t test. *P* < 0.05 were considered statistically significant.

Due to the ethical requirement for experimental animals, we had to stop the experiment at day 69 because the kidneys from control mice displayed severe pathological damages. However, the circUOX-LNP–treated mice exhibited markedly alleviated kidney lesions (Fig. 5B and Fig. S4). In addition, no mortality was observed in the treatment group during the study, in contrast to several deaths recorded in the control cohort (Fig. 5C). Moreover, circUOX-LNP reduced serum urea and creatinine levels, indicating the alleviated renal damage in treatment group (Fig. 5D).

Meanwhile, serum liver function indicators (ALT, AST and ALP) and histological examination of hearts, lungs, spleens, and livers all suggested no detectable toxicity from long-term circUOX-LNP treatment. (Fig. S5A and Fig S5B). Collectively, these results demonstrate that the weekly administration of circUOX-LNP provides sustained urate-lowering efficacy and effectively mitigates renal injury in the hyperuricemia mouse model.

## Discussion

This study presents the first proof-of-concept for the treatment of severe hyperuricemia using circRNA-based therapy, showcasing the long-term efficacy of circRNA-mediated uricase replacement. Recently, the mRNA encoding mouse uricase was used to reduce serum urate levels in an asymptomatic hyperuricemia mouse model generated by a high-purine diet and siRNA-mediated knockdown of mouse uricase (*60*). However, the efficacy of such treatment was modest, and the therapeutic effect of repeated dosing was not evaluated (*60*). In contrast, we employed a UOX-knockout model to mimic human uricase deficiency (*55*), which shows severe hyperuricemia and develops renal injury at early age. This model is particularly suitable for studying refractory hyperuricemia. In addition, we used a circRNA encoding humanized uricase, which showed high enzymatic activity and efficient peroxisomal targeting in human cells. We found that a single injection of circUOX-LNP in diseased mice rapidly reduced serum urate within 24 h, and the repeated dosing generated a prolonged urate-lowering effect that effectively protected against hyperuricemia-induced renal damage.

This circRNA-based therapy will potentially be beneficial to patients who are ineligible for first-line drugs, such as those with allopurinol hypersensitivity or pegloticase resistance. The rapid urate reduction achieved by a single injection in mouse model also highlights its potential advantages over xanthine oxidase inhibitors in conditions requiring immediate intervention, such as tumor lysis syndrome. Moreover, the sustained efficacy over a 10-week dosing regimen is particularly significant for clinical translation, as severe primary hyperuricemia is typically incurable and requires lifelong management. Considering that urate levels are influenced by renal excretion efficiency, and that UOX^⁻/⁻^ mice exhibit more severe hyperuricemia and accelerated renal injury compared with humans, we anticipate that the effective dose required in clinical applications is likely to be lower, which may favor a long-term treatment. Furthermore, circRNAs may also offer advantages over current linear mRNA drugs because it can bypass the need for extensive nucleoside modifications that is known to cause translation frameshift (*61*).

In this study, we showed that circRNA-LNPs exhibited a favorable biosafety profile with sustained efficacy in repeated dosing. However, it remains unclear if such repeat dosing is applicable for life-long treatment in humans, as the low toxicity observed in mice may not be transferable to humans. Therefore, further optimization should focus on engineering IRES and circRNA architectures that could enhance translation efficiency and extend expression duration in hepatocytes, thereby reducing the dosing frequency, minimizing potential toxicity and extending the therapeutic window. Similarly, improvements in delivery vehicles may also help to enhance hepatic targeting and further extend protein expression. Moreover, the development of engineered uricase variants with improved catalytic activity and extended half-life represents another promising strategy to increase therapeutic potency.

In conclusion, circRNA-based uricase replacement combines rapid urate reduction with sustained long-term efficacy, establishing a promising strategy to address unmet clinical needs in hyperuricemia management. The concept of repeated circRNA administration for sustained expression of therapeutic proteins in the liver may extend beyond uricase replacement, providing a therapeutic strategy for other severe protein-deficiency disorders that require long-term treatment.

## Materials and Methods

### Study Design

Three candidate uricase sequences (rHU, PBC, and AN96) were synthesized as circRNAs to assess expression and uricase activity in human cells. Human 293T and HepG2 cells were transfected, and uricase expression and activity of cell lysates were evaluated by western blotting and uricase activity assays (*n* = 3 per group). Based on superior enzymatic activity, uricase circRNA rHU was selected for peroxisomal targeting verification and subsequent in vivo studies. Lipid nanoparticles were encapsulated to deliver circRNAs to mouse livers via tail vein injections.

To determine the efficiency of liver-targeted circRNA delivery and potential hepatic or renal toxicity, wild-type C57BL/6J mice (*n* = 3 per group) were injected via the tail vein with varying doses of circUOX-LNPs (0.3, 0.6, or 1.2 mg/kg). Mice were sacrificed on days 1, 3, and 5 for collection of serum and major organs. Untreated mice served as the healthy group, and mice injected with Tris-HCl (PH=7.5, 50mM) served as the control group and were sacrificed at day 1. Hepatic circRNA expression was quantified by RT-qPCR and western blotting, and serum indicators of liver and kidney function (ALT, AST, ALP, ALB, urea, and creatinine) were measured to evaluate biosafety.

To evaluate the efficacy of circUOX-LNP in hyperuricemia, circUOX-LNPs were intravenously injected to uricase knockout mice (*n* = 6-8 per group), with Tris-HCl buffer serving as the control. Serum urate levels were measured at multiple time points before and after circUOX-LNP administration via retro-orbital blood collection. Serum indicators were assessed for biosafety profile and major organs were H&E stained to detect organ damages. For single-dose experiments, mice were monitored until serum urate returned to baseline. To evaluate the efficacy of multiple doses, circUOX-LNPs were administered every four days based on the duration of the single-dose effect, and mice were sacrificed eight days after the third injection. Three mice were orally administrated with allopurinol every day as a positive control. To assess long-term efficacy, circUOX-LNPs were administered every eight days for up to 69 days; the experiment was terminated due to severe health deterioration of control group.

### CircRNA synthesis and LNP production

The protein coding fragments of UOX rHU, PBC, and An96 were chemically synthesized from Genewiz (Suzhou) Inc. and cloned by Vazyme™ Cloning Kit (Vazyme, c112) into the plasmid backbone containing T7 RNA polymerase promoter, 5’ group II self-splicing intron, CVB3 IRES, FLAG tag and 3’ group II self-splicing intron. UOX circRNA was synthesized from the linearized DNA template by in-vitro transcription (IVT) with T7 RiboMAX™ Large-Scale RNA Production System (Promega, P1300). CircRNA was purified with the help of Shanghai CirCode Biomedicine Inc. using the high performance liquid chromatography (HPLC) system. Sequences for IVT plasmid elements were listed in Supplementary Data 1.

The circRNA was encapsulated in lipid nanoparticles using a lipid formulation via the NanoAssemblr Ignite system with the support of Shanghai CirCode Biomedicine Inc. Lipid mixture and aqueous phase containing circRNA were mixed by microfluidics. After that, circRNA-LNP was ultra-filtrated, concentrated to the desired level, and stored at −80 °C in a 50mM Tris-HCL buffer (PH = 7.5) containing 10% sucrose. The formulation of the lipid mixture is ALC-0315:DSPC:Cholesterol:ALC-0159 = 46.3:9.4:42.7:1.6.

### Mammalian cell culture and transfection

293T and HepG2 cells were cultured in DMEM media containing 10% FBS at 37 °C supplied with 5% CO2. Cells were seeded on 6-well plates or 12-well plates one day before transfection. CircRNAs were transfected with Lipofectamine MessengerMAX™ (Themofisher, LMRNA001) according to the manufacturer’s protocol. Transfected cells were collected 48 h after transfection for further RNA and protein analysis.

### RT-qPCR

Total RNAs were isolated using TRIzol™ Reagent from cells according to the manufacturer’s protocol. 1μg total RNA was reverse-transcribed with ABScript™ Neo RT Master Mix (Abclonal, RK20433). The real-time PCR was performed using Hieff™ qPCR SYBR Green Master Mix (Yeasen, 11201ES03) and BIOER LineGene™ 9600 Plus Real-time qPCR System according to the manufacturer’s instructions. All the primers used in qPCR were listed in Supplementary Data 2.

### Western blot

The cell lysates were mixed with SDS loading buffer (Beyotime, P0015), heated at 100 °C for 10 min, separated by SDS-PAGE gels and transferred to PVDF membranes. Membranes were blocked in 5% milk for 30 minutes and incubated overnight at 4°C with the following antibodies: flag tag antibody (Cell Signaling Technology, 14793) and alpha-tubulin antibody (Proteintech, HRP-66031). HRP-linked secondary antibodies were used and blots were visualized with the ECL Kit (NCMBiotech, P10100) in the Bio-Rad imaging system. The original Western blot images are provided in Supplementary Data 3.

### Uricase activity assay with cell lysates

Cells were lysed in moderate lysis buffer (Beyotime, P0013) with protease inhibitor cocktail. Total protein concentrations of cell lysates were measured by the BCA method. The uricase enzymatic activity of cell lysates was measured in 0.1 M sodium borate buffer (pH 8.5) with 0.125mM uric acid at 37°C for 120 min by the decrease in absorbance at 293 nm due to enzymatic oxidation of uric acid. Uric acid (Sigma-Aldrich, U2625) was used to prepare standards. Recombinant uricase (0.02µg, purchased from Beijing BIOSS Biotech Inc., D10166) was used as a positive control.

### Measurement for the co-localization of UOX and peroxisomes

GFP coding sequence was fused to N-terminal of rHU sequence to labeled rHU *in vivo*, while a peroxisome targeting signal sequence SRL (Ser-Arg-Leu) (*62*) was added to C-termical of mCherry to labeled peroxisomes. 293T cells were co-transfected with Lipofectamine™ 3000 (Thermofisher, L3000001). Images of living cells were captured with a 60x oil immersion objective by using the Olympus FV1200 Laser Scanning Confocal Microscope.

### Animal care and maintenance

The wild type C57BL/6J and UOX-knockout mice were purchased form Cyagen Biosciences (Suzhou) Inc. and were maintained under specific pathogen-free conditions, a 12-h light/12-h dark cycle and temperatures of 20-26 °C with 40-70% humidity. All mouse experiments were conducted in accordance with protocols approved by the Shanghai Institute of Nutrition and Health, CAS.

### Pharmacokinetics and biosafety assay in wild type mice

Wild type 7-week-old male C57BL/6J mice received circUOX-LNP via tail vein at 0.3, 0.6, or 1.2 mg/kg. Mice injected with each dose were sacrificed at day 1, 3 and 5 post injection (*n* = 3 per group). Mice of control group were injected with Tris-HCl (PH = 7.5, 50mM) buffer solution and were sacrificed at day 1 (*n* = 3). Untreated mice were used as the healthy group (*n* = 3). Major organs and serums were collected for further analysis to detect quantities of circRNA and proteins, as well as other biochemical indicators for liver and kidney functions.

To detect uricase protein in mouse liver, we concentrate the circRNA encoded uricase using anti-FLAG immunomagnetic beads to avoid high non-specific background from liver lysates. The MagBeads (Yeason, 20565ES03) was used according to manufacturer’s instructions to Immunoprecipitate FLAG-tagged uricase expressed by exogenous UOX circRNA in mice hepatic lysates. Briefly, 10ul magnetic beads were incubated with hepatic lysates of 5mg total protein overnight at 4°C with gentle rotation. Beads were then washed three times with lysis buffer to remove nonspecific binding proteins. Bound proteins were eluted by boiling the beads in 50 μl SDS loading buffer for 10 min, 20 μl of which was used in the Western blot analysis.

The analysis of serum biochemical indicators including ALT, AST, ALP, ALB, urea, and creatinine were performed by the automatic biochemical analyzer in Wuhan Boerfu Biotechnology Co., Ltd.

### Efficacy study in UOX*^-/-^* hyperuricemia mouse model

UOX knockout mice were purchased and retained in Cyagen Biosciences. UOX^-/-^ mice were maintained with water containing allopurinol (90mg/L) after birth until 2-5 days before experiments. In the single-injection study, 6-week-old mice (*n* = 8 per group) were employed and water with allopurinol were replaced with normal water 5 days before the injection. CircUOX-LNP was injected via tail vein at 0.6 or 1.2 mg/kg. An equal volume of buffer solution (50 mM Tris-HCl, PH = 7.5) was injected as control group. Serums were collected at 1, 3, 5, 7, 10, 15 days after injection. In the three-injection study, 8-week-old mice (*n* = 7 per group) were used and water was replaced 5 days before the first injection. CircUOX-LNP of 1.2mg/kg or an equal volume of buffer was injected via tail vein at day 0, day 4 and day 8. Three mice were orally administrated with 0.3mg allopurinol every day as a positive control. Serums were collected at 1, 3, 5, 7, 10, 13, 15, 20 days after the first administration. In the long-term study of 69 days, 4-week-old mice (*n* = 6 per group) were used and water with allopurinol was replaced 2 days before the experiment. The FLAG-tag was removed from circUOX sequence. CircUOX-LNP of 1.2mg/kg or an equal volume of buffer solution was injected every 8 days after the serum collections. Serums were collected at 2, 5, 8 days after each administration. Mice were injected total 8 times. Mice in all three studies were sacrificed at the end of experiments and hearts, livers, spleens, lungs, kidneys, and serums were collected for further analysis.

### Detection of serum uric acid

Mouse blood was obtained from the eye socket and stand at room temperature for 60 min. After centrifugation at 1000 × g for 15 min, the supernatant was collected as serum. The concentration of serum uric acid was measured using Amplex™ Red Uric Acid/Uricase Assay Kit (Thermofisher, A22181) following the manufacturer’s instructions.

### Hematoxylin and eosin (H&E) staining

Mouse tissues were fixed in 4% paraformaldehyde at room temperature for 24 h, dehydrated, embedded in paraffin and sectioned to 5 μm. Sections were deparaffinized, rehydrated, and stained with hematoxylin and eosin and examined using the Pannoramic MIDI digital scanner (3DHISTECH Ltd.) in Wuhan Boerfu Biotechnology Co., Ltd. following standard protocols.

### RNA seq

RNA-seq was accomplished by Genewiz (Suzhou) Inc. Briefly, total RNA was extracted and assessed for quality. 1 μg of total RNA was used for library preparation. mRNA was enriched using Oligo(dT) beads, fragmented, and reverse-transcribed into cDNA. Libraries were constructed, amplified, and sequenced on an Illumina platform to generate paired-end reads. Raw sequencing reads were trimmed to remove adapters and low-quality bases using Cutadapt (v1.9.1) and quality-checked using FastQC. Clean reads were aligned to the reference genome using HISAT2 (v2.2.1), and gene-level counts were quantified using HTSeq (v0.6.1). Differential expression analysis was performed using DESeq2 (v1.34.0), with significance defined as |Foldchange value| >1.5 and adjusted p-value < 0.05. Functional enrichment analyses were conducted using Metascape analysis. RNA-seq data generated in this study have been deposited in the Gene Expression Omnibus (GEO) under accession number GSE312701.

### Statistical analysis

All data were presented as mean ± SD. Statistical significance between groups for uricase activity, relative mRNA expression or serum indicators was assessed using unpaired t-test or one-way ANOVA followed by Dunnett’s multiple comparisons test. The significance of urate levels in the efficacy study was assessed by two-way ANOVA followed by Bonferroni’s multiple comparisons test. All analysis used GraphPad Prism 9.0.0 software. *P* values less than 0.05 were considered statistically significant (*ns* > 0.05, \**p* < 0.05, \*\**p* < 0.01, \*\*\**p* < 0.001, \*\*\*\**p* < 0.0001).

## Author contributions

Conceptualization, Z.W.; methodology, Z.Z., J.Z., K.Z., J.H., Y.Y., and Z.W.; validation, Z.Z and Z.W.; formal analysis, Z.Z. and J.Z.; investigation, Z.Z.; resources, Z.Z., J.Z., K.Z., J.H., Y.Y., and Z.W; data curation, Z.Z and J.Z; writing - original draft preparation, Z.Z.; writing - review and editing, Z.Z., J.Z. and Z.W.; supervision, Y.Y and Z.W.; project administration, Z.W.; funding acquisition, Z.W.; Y.Y. and the CirCode BioMed team (K.Z, J.H) produced circRNAs and performed quality control of circUOX-LNP in the animal work. All authors have read and agreed to the published version of the manuscript.

## Acknowledgements

We thank Ms. Yun Jiang for her assistance in preparing the manuscript, and the members of the Wang laboratory for their helpful discussions and comments. This work is supported by National Natural Science Foundation of China (32030064 and 32250013 to Z.W.), the National Key Research and Development Program of China (2021YFA1300503 to Z.W), and the internal fund of Shanghai CirCode Biomed.

## Supplementary Materials

### Supplementary Figures and Figure Legends

**Figure.S1.**
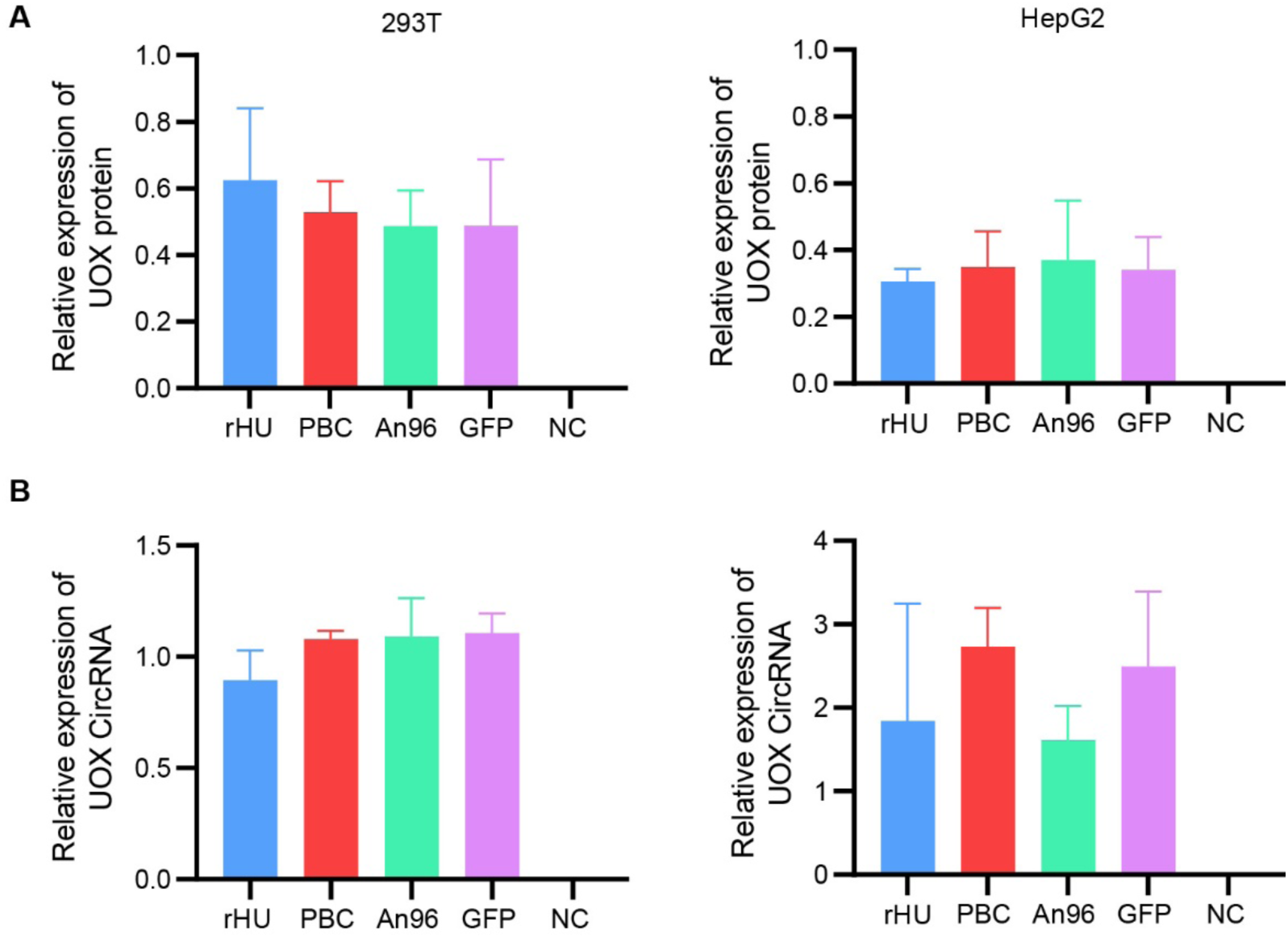
Relative quantification of UOX circRNA and protein in transfected cells. (A) Relative quantification of UOX expression in western blot in 293T and HepG2 cells transfected with rHU, PBC, An96 and GFP circRNAs, normalized to α-tubulin. (B) Relative levels of UOX circRNAs in 293T and HepG2 cells transfected with rHU, PBC, An96 and GFP circRNAs. RT-qPCR was performed using primers flanking the back-splicing junction, and circRNA levels were normalized to GAPDH mRNA levels. Data were presented as mean ± SD.

**Figure.S2.**
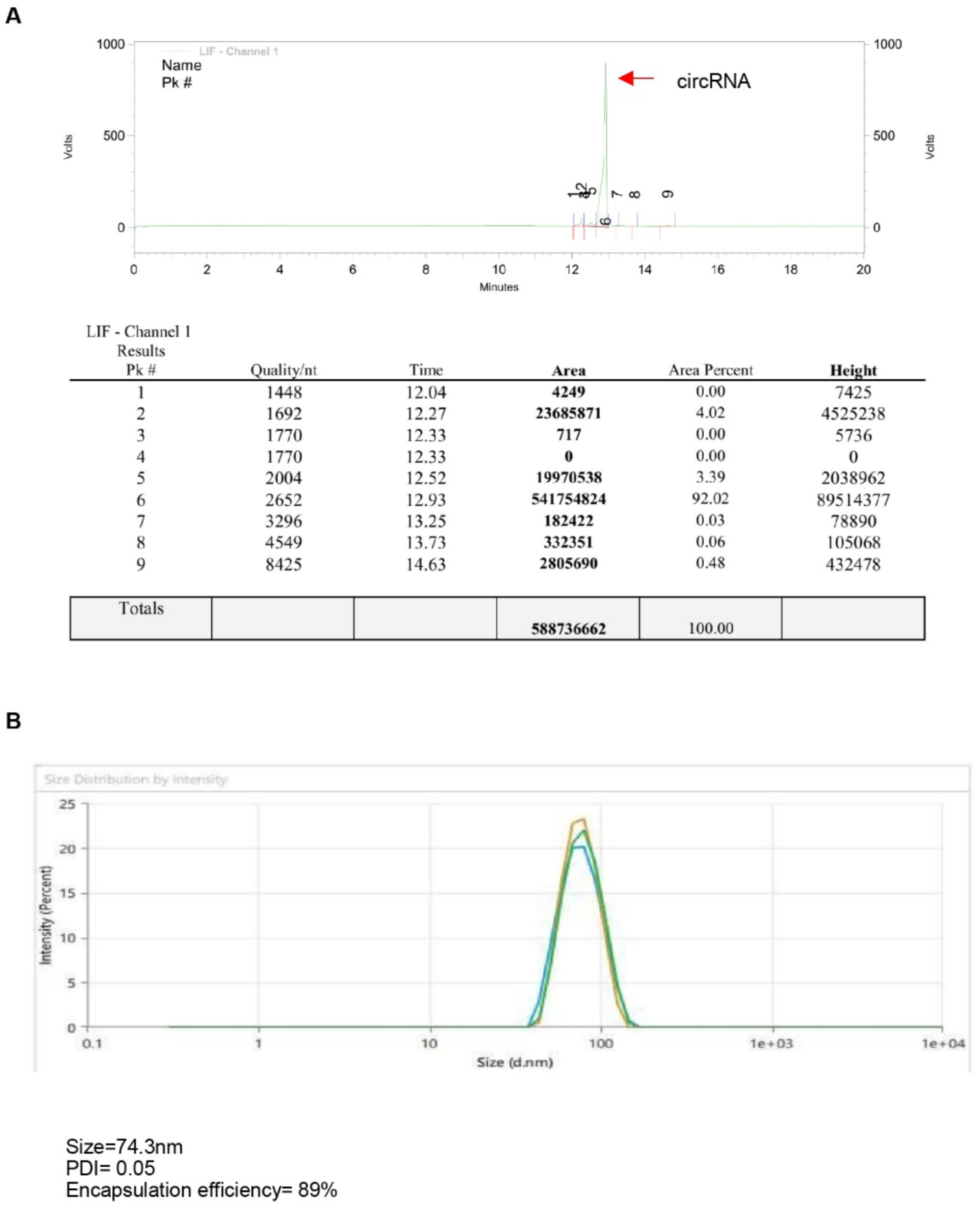
Characterization of rHU circRNA and circRNA-LNPs. (A) Capillary electrophoresis of purified the UOX rHU circRNA. The peaks were automatically assigned, and the peak 6 corresponding to circRNA. (B) Particle size analysis of UOX rHU circRNA-LNPs.

**Figure S3.**
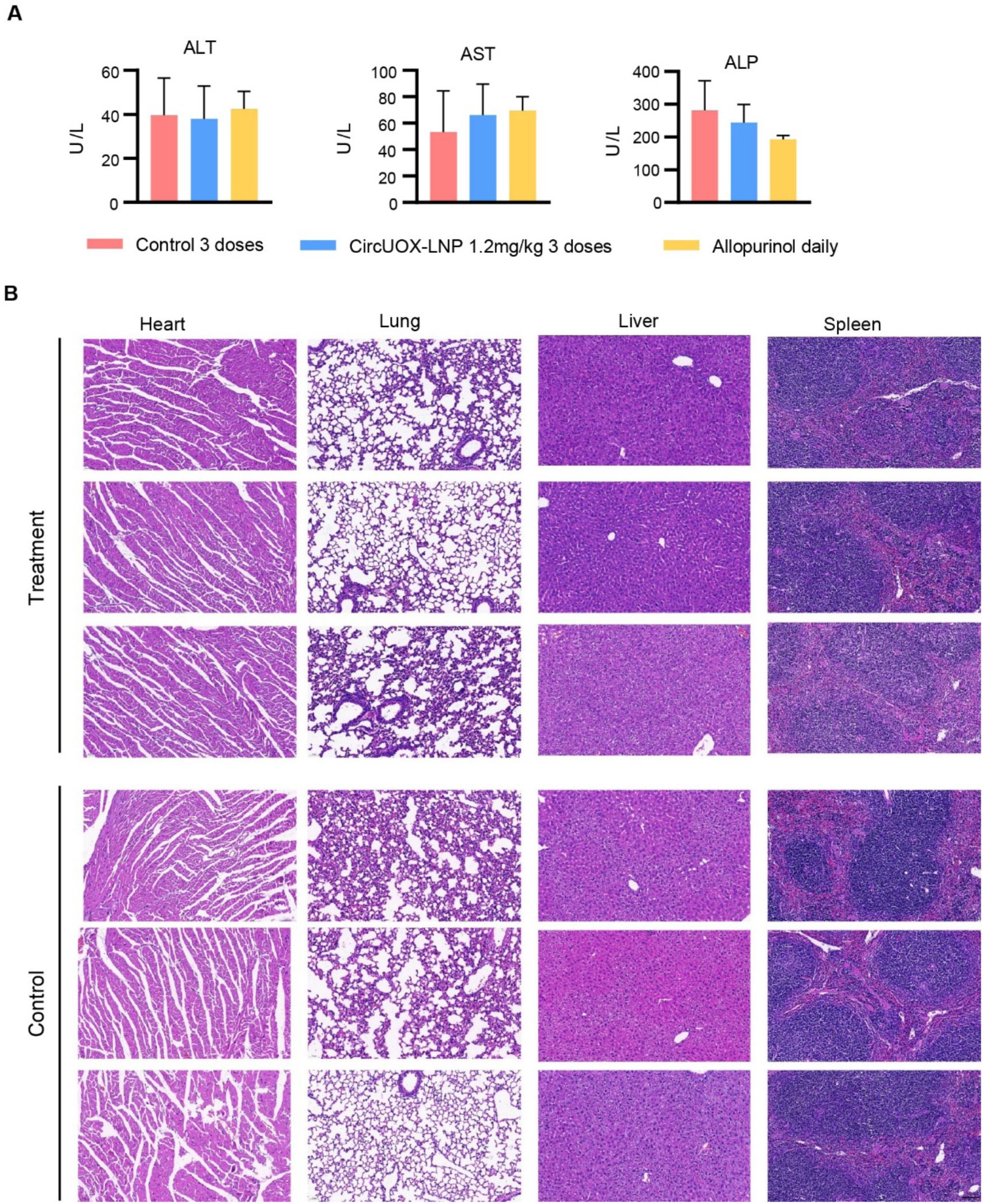
Biosafety of three-dose circUOX-LNP. (A). Serum hepatic function indicators (ALT, AST, ALP) in the UOX^-/-^ mice after three doses of 1.2mg/kg circUOX-LNP. Mice receiving three doses of buffer or daily allopurinol gavage served as controls. Five serum samples were analyzed per group. (B) H&E staining of hearts, livers, spleens, and lungs from the UOX^-/-^ mice receiving 3 doses of circUOX-LNP or buffer. Three organ samples were examed per group. Scale bar, 200µm.

**Figure S4.**
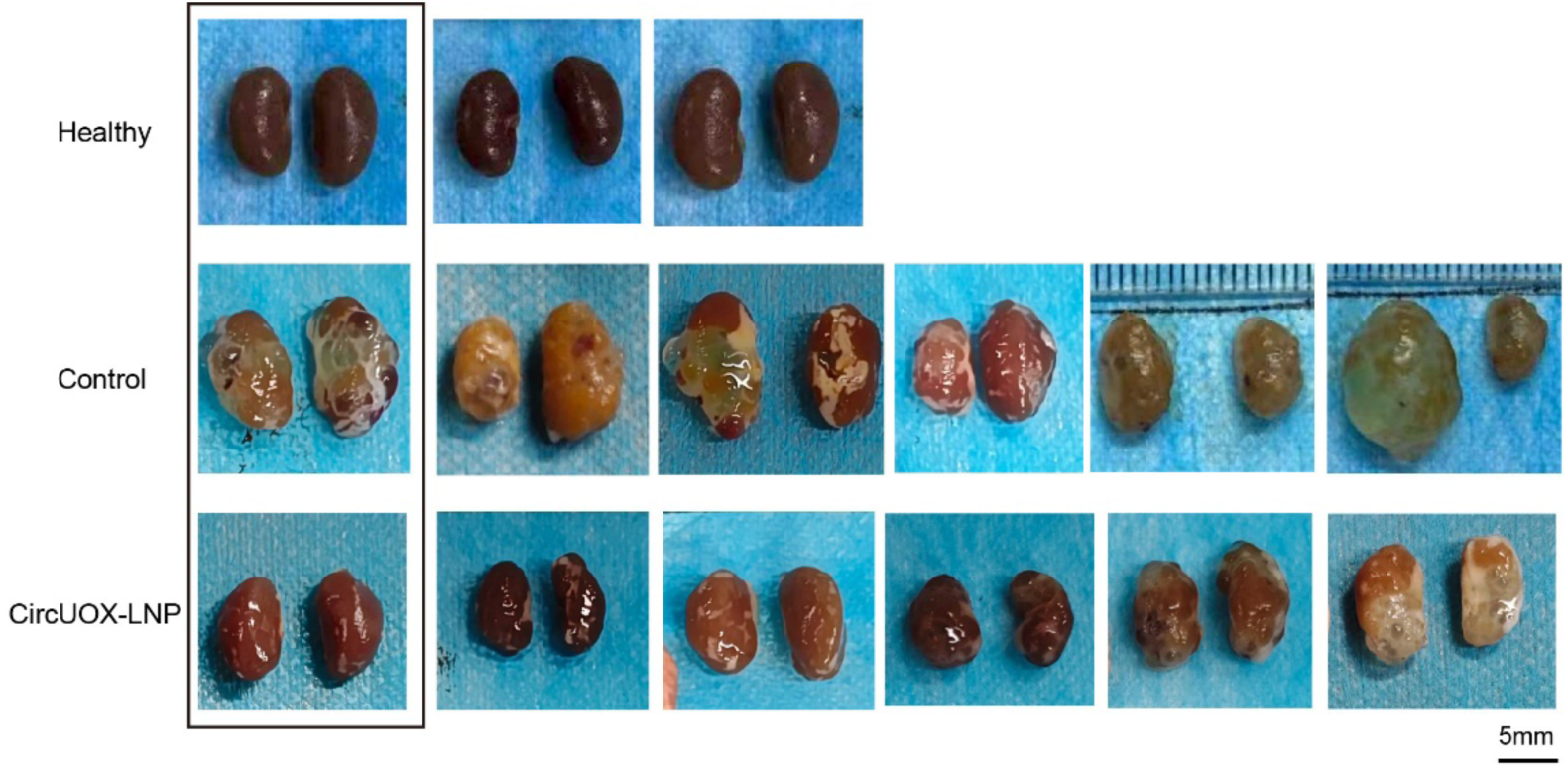
Kidneys of healthy mice, control group and treatment group in the long-term efficacy study. The UOX^⁻/⁻^ mice receiving eight injections of 1.2mg/kg circUOX-LNP or buffer served as the treatment and control groups (*n* = 6), whereas wild type mouse served as the healthy group (*n* = 3). CircUOX-LNPs were administered once every eight days. The mice were sacrificed on day 69 and all kidney images were shown. The images of first columns were used in Fig. 5B. Scale bar, 5mm.

**Figure S5.**
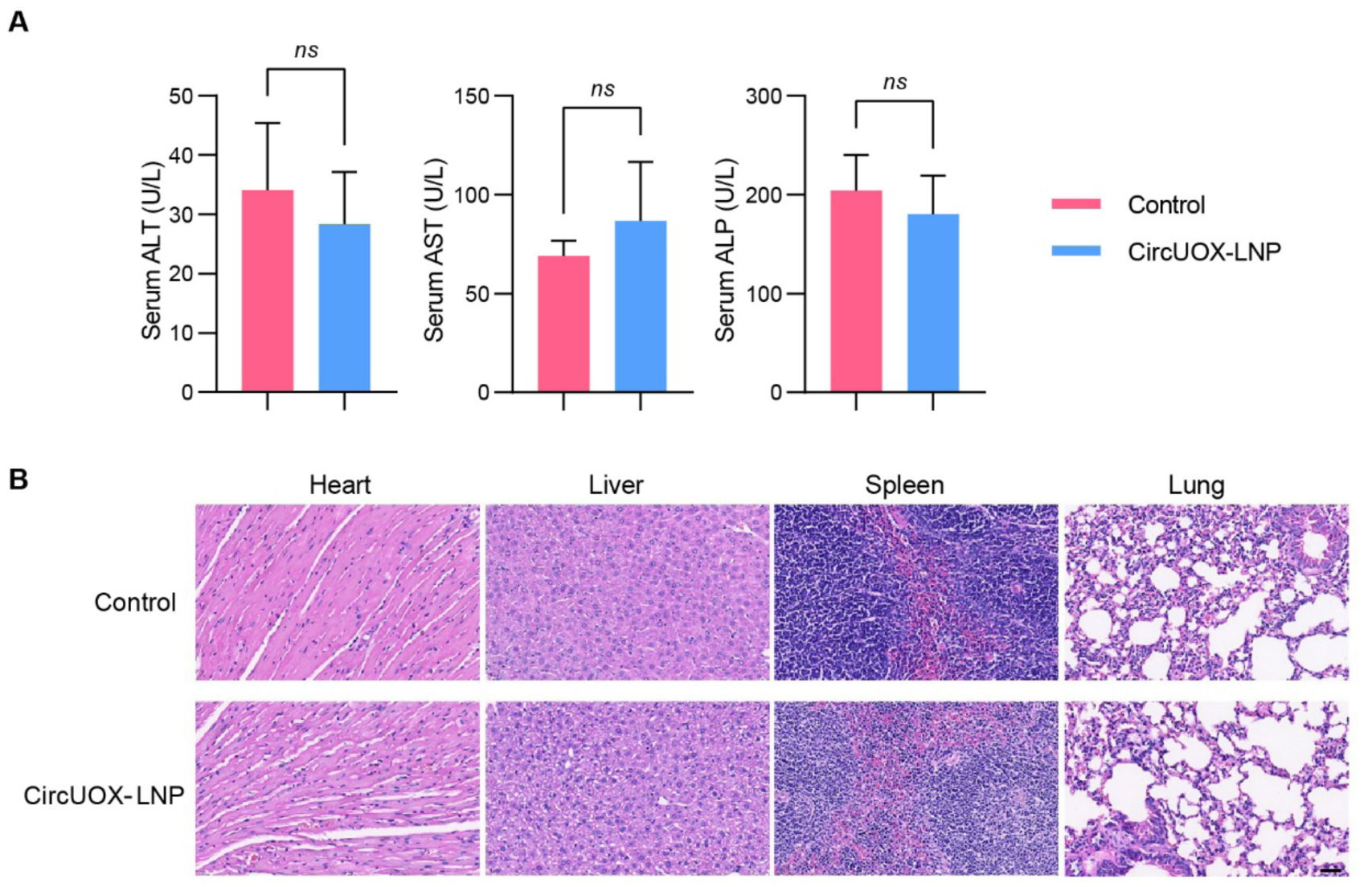
Biosafety of long-term circUOX-LNP administration in UOX*^-/-^*mice. (A) Serum hepatic function indicators (ALT, AST, ALP) in the UOX^-/-^ mice recieving eight doses of 1.2mg/kg circUOX-LNP or buffer (*n* = 6). Data were presented as mean ± SD. Statistical significance was determined by unpaired t-test between circUOX-LNP treated group and control group. *p* > 0.05 was considered no statistically significant (*ns*). (B) H&E staining of hearts, livers, spleens, and lungs from the UOX^-/-^ mice treated with 8 doses of circUOX-LNP or buffer. Three organ samples were examined per group. Scale bar, 40µm.

